# Replication of [AT/TA]_25_ microsatellite sequences by human DNA polymerase δ holoenzymes is dependent on dNTP and RPA levels

**DOI:** 10.1101/2023.11.07.566133

**Authors:** Kara G. Pytko, Rachel L. Dannenberg, Kristin A. Eckert, Mark Hedglin

## Abstract

Difficult-to-Replicate Sequences (DiToRS) are natural impediments in the human genome that inhibit DNA replication under endogenous replication. Some of the most widely-studied DiToRS are A+T-rich, high “flexibility regions,” including long stretches of perfect [AT/TA] microsatellite repeats that have the potential to collapse into hairpin structures when in single-stranded DNA (ssDNA) form and are sites of recurrent structural variation and double-stranded DNA (dsDNA) breaks. Currently, it is unclear how these flexibility regions impact DNA replication, greatly limiting our fundamental understanding of human genome stability. To investigate replication through flexibility regions, we utilized FRET to characterize the effects of the major ssDNA-binding complex, RPA, on the structure of perfect [AT/TA]_25_ microsatellite repeats and also re-constituted human lagging strand replication to quantitatively characterize initial encounters of pol δ holoenzymes with A+T-rich DNA template sequences. The results indicate that [AT/TA]_25_ sequences adopt hairpin structures that are unwound by RPA and pol δ holoenzymes support dNTP incorporation through the [AT/TA]_25_ sequences as well as an A+T-rich, non-structure forming sequence. Furthermore, the extent of dNTP incorporation is dependent on the sequence of the DNA template and the concentration of dNTPs. Importantly, the effects of RPA on the replication of [AT/TA]_25_ sequences are dependent on the concentration of dNTPs, whereas the effects of RPA on the replication of an A+T-rich, non-structure forming sequence are independent of dNTP concentration. Collectively, these results reveal complexities in lagging strand replication and provide novel insights into how flexibility regions contribute to genome instability.

## Introduction

Human genome replication requires intimate coordination of the diverse enzyme activities that comprise the DNA replication machinery (i.e., replisome) as well as the maintenance of abundant metabolite pools. Loss of coordinated replisome function and/or depletion of metabolite pools from endogenous or exogenous sources leads to DNA replication dysfunction (i.e., replication stress) that can ultimately generate DNA replication errors, DNA strand breaks, and genome rearrangements. These alterations of the human genome are collectively referred to as genome instability and are all hallmarks of carcinogenesis (1).

Difficult-to-Replicate Sequences (DiToRS) are natural impediments in the human genome that inhibit DNA replication under endogenous replication stress conditions (2). Some of the most widely studied DiToRS are within chromosomal fragile sites, which are unstable genomic regions prone to gap formation or breakage during DNA replication (3). Several common fragile sites (CFSs) contain A+T-rich, high “flexibility regions,” that include stretches of [A/T] and [AT/TA] microsatellite repeats, the latter of which have the potential to collapse into hairpin structures when in single-stranded DNA (ssDNA) form (4-7). Genome-wide analyses revealed that A+T-rich CFS sequences prone to hairpin formation coincide with chromosomal hotspots for genome re-arrangements (8-14). While many factors contribute to CFS instability such as chromatin structure, replication origins, and replication-transcription conflicts (15,16), direct evidence exists for DNA replication stalling within specific A+T-rich CFS regions (3,13,17-20). In particular, recent studies from the Nussenzweig lab observed that expanded [TA]_n_ repeats form intramolecular DNA secondary structures and are preferential sites for DNA replication stalling and replication fork collapse (21-23). Currently, the mechanisms by which replication stress causes DNA replication stalling at/within [AT/TA] microsatellite repeats are unknown.

In humans, like all eukaryotes, lagging strand DNA templates are immediately engaged by the major ssDNA-binding complex, replication protein A (RPA), as they are exposed and then replicated primarily by DNA polymerase δ (pol δ) holoenzymes comprised of pol δ and the processivity sliding clamp, proliferating cell nuclear antigen (PCNA) (24-27). Hence, A+T-rich, “high flexibility” regions in lagging strand DNA templates are initially encountered by pol δ holoenzymes during scheduled DNA replication in S-phase. To investigate how replication stress causes DNA replication stalling at/within [AT/TA] microsatellite repeats, we characterized the structures of A+T-rich DNA sequences from human CFSs and their replication by pol δ holoenzymes. Specifically, [AT/TA]_25_ microsatellite repeats were directly compared to an A+T-rich DNA sequence that is devoid of any repetitive elements. Förster Resonance Energy Transfer (FRET) analyses reveal that [AT/TA]_25_ microsatellite repeats adopt hairpin structures that are rapidly unwound by abundant RPA. Quantitative biochemical assays that analyze initial encounters of human pol δ holoenzymes with DNA sequences reveal that pol δ holoenzymes support dNTP incorporation through both the [AT/TA]_25_ microsatellite repeat and the A+T-rich, non-structure-forming sequence. The extent of dNTP incorporation is dependent on the template sequence and the concentration of dNTPs. Importantly, our studies also reveal that the effects of RPA on the replication of [AT/TA]_25_ microsatellite repeats are dependent on the concentration of dNTPs, whereas the effects of RPA on the replication of an A+T-rich, non-structure forming sequence are independent of dNTP concentration. Collectively, the results from the present study reveal complexity in the structural dynamics and replication of lagging strand DNA templates and provide novel insights into how flexibility regions contribute to genome instability.

## MATERIALS AND METHODS

### Recombinant Human Proteins

Human RPA, PCNA, Cy5-PCNA, RFC, and exonuclease-deficient pol δ were obtained as previously described (28,29). Herein, exonuclease-deficient pol δ is simply referred to as pol δ. The concentration of active RPA was determined via a FRET-based activity assay, as described previously (30).

### Oligonucleotides

A+T-rich DNA templates to be replicated by pol δ (FRA3B-AT3, FRA3B-AT3-RC, FRA16D-C, **Table 1**) are incorporated into the analytical procedures described below by distinct modifications (depicted in **Figure S1**). The control DNA template (FRA16D-C) has been previously described (31) and is in a region of the human FRA16D CFS that is 76 % A+T but devoid of any repetitive elements. The DNA templates containing an [AT/TA]_25_ repeat (FRA3B-AT3 and FRA3B-AT3-RC) have been previously described (32) and are within the AT3 region of the human FRA3B CFS that is 80 % A+T. FRA3B-AT3-RC is the reverse complement of FRA3B-AT3 (32-35).

**Table 1:**
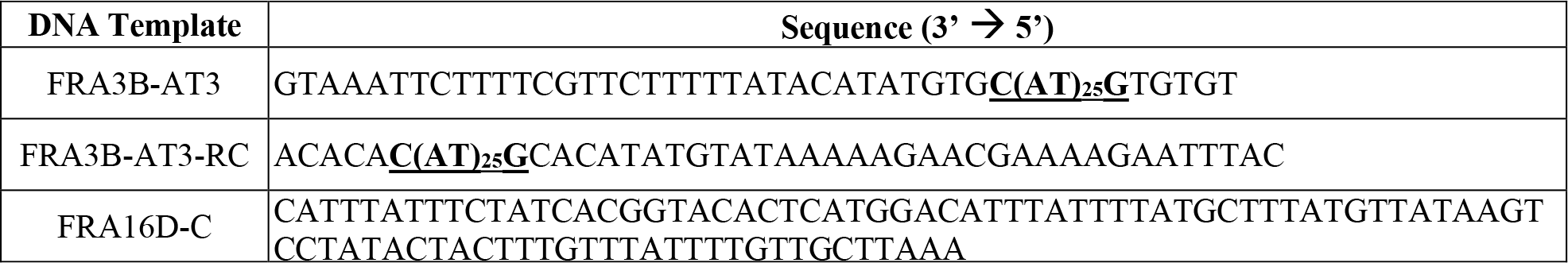
A+T-rich DNA template sequences analyzed in this study. Sequences are written in the direction (3’ → 5’) in which they are interpreted by a pol δ holoenzyme. Potential hairpin-forming sequences in FRA3B-AT3 and FRA3B-RC are in bold and underlined.

Oligonucleotides were synthesized by Integrated DNA Technologies (Coralville, IA) or Bio-Synthesis (Lewisville, TX) and purified on denaturing polyacrylamide gels. The concentrations of unlabeled DNAs were determined from the absorbance at 260 nm using the provided extinction coefficients. The concentrations of Cy5-labeled DNAs (including those that contain a Cy3 label) were determined from the extinction coefficient at 650 nm for Cy5 (ε_650_ = 250,000 M^−1^cm^−1^). For annealing two ssDNAs (as depicted in **Figure S1**), the primer and corresponding complementary template strand were mixed in equimolar amounts in 1X Annealing Buffer (10 mM TrisHCl, pH 8.0, 100 mM NaCl, 1 mM EDTA), heated to 95 ^o^C for 5 min and allowed to slowly cool to room temperature. For annealing hairpin DNAs (as depicted in **Figure 1** and **S1**), the DNA sequence (1 μM in 1X Annealing Buffer) was heated to 95 ^o^C for 5 min and then immediately submerged in ice water.

**Figure 1.**
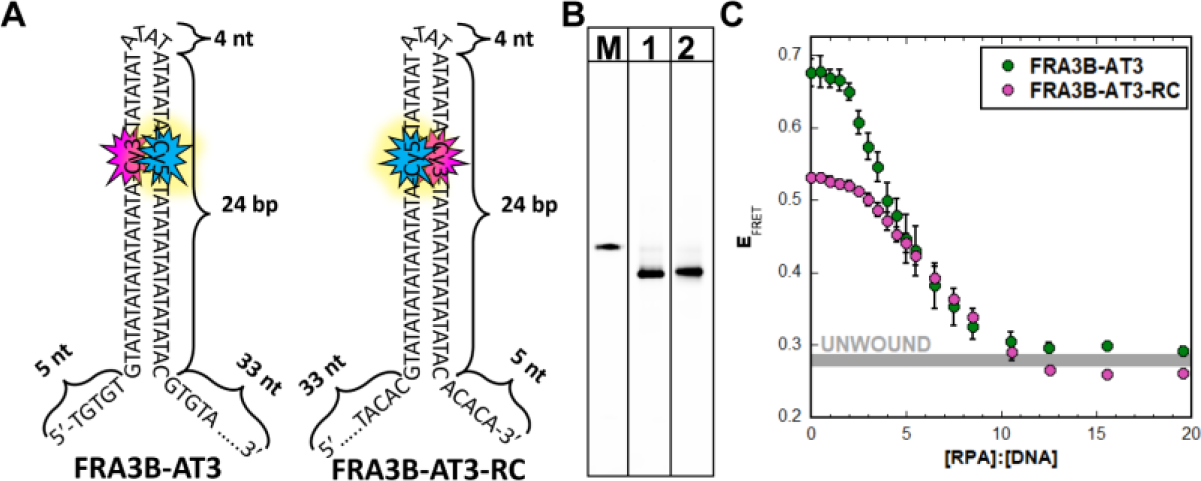
[AT/TA]_25_ sequences form hairpins that are unwound by RPA. (**A**) Schematic representation of the Cy3/Cy5-labeled DNA sequences. When annealed as hairpins (as depicted), the Cy3 FRET donors and Cy5 FRET acceptors are juxtaposed. (**B**) Native gel electrophoresis of oligonucleotides. A blunt-ended 26 bp duplex with 33 nt (5′) and 5 nt (3′) ssDNA overhangs at the opposite end (**Figure S1**) was run on the gel as a size marker (lane M) for the FRA3B-AT3 (lane 1) and FRA3B-AT3-RC (lane 2) hairpins. (**C**) Effect of RPA on the structure of [AT/TA]_25_ sequences. FRA3B-AT3 or FRA3B-AT3-RC hairpins were titrated with RPA and the equilibrium E_FRET_ values were monitored. Each data point represents the average + S.E.M. of at least three independent experiments. The equilibrium E_FRET_ values (± S.E.M.) observed for complete unwinding (“UNWOUND,” determined in **Figure S2**) is indicated as a horizontal grey bar. Results for each Cy3/Cy5-labeled hairpin are plotted as a function of the respective [RPA]:[DNA] ratios.

### Native Gel Electrophoresis

Annealed DNA substrates were diluted in 1X Replication Buffer [25 mM HEPES•OAc, pH 7.5, 10 mM Mg(OAc)2, 125 mM KOAc] supplemented with 0.1 mg/ml BSA and 1 mM TCEP, and the final ionic strength was adjusted to physiological (200 mM) by the addition of appropriate amounts of KOAc. Samples were run on 15 % native PAGE (19:1 acrylamide:bisacrylamide, 0.5 X TBE) at constant voltage of 10 v/cm. Gels images were obtained on a Typhoon Model 9410 fluorescence imager (GE Healthcare).

### FRET Measurements

All experiments were performed at room temperature (23 ± 2 °C) in 1X Replication Buffer supplemented with 1 mM DTT, and the final ionic strength was adjusted to physiological (200 mM) by the addition of appropriate amounts of KOAc. All measurements were performed in 16.100F-Q-10/Z15 sub-micro fluorometer cells (Starna Cells) and analyzed in a Horiba Scientific Duetta-Bio fluorescence/absorbance spectrometer. Excitation and emission slit widths were each set to 5 nm, unless indicated otherwise. Reaction solutions were excited at 514 nm and the fluorescence emission intensities (*I*) were simultaneously monitored at 563 nm (*I*_563_, Cy3 FRET donor fluorescence emission maximum) and 665 nm (*I*_665_, Cy5 FRET acceptor fluorescence emission maximum) over time, recording *I* every 0.17 s (36). For each time point, E_FRET_ was calculated where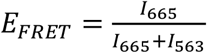. For RPA titrations, a solution containing 20 pmol of a Cy3/Cy5-labeled hairpin (**Figure S1**) was titrated with increasing amounts of RPA (heterotrimer) and the equilibrium E_FRET_ value at each amount of RPA was calculated as follows. E_FRET_ values were recorded every 0.17 s beginning 10 s after the addition of RPA and continued until the E_FRET_ maintained a constant value for at least 1 min. All E_FRET_ data points within this stable region were then averaged to obtain the equilibrium E_FRET_ value for a given RPA amount (37,38). For all plots in all figures, each data point/column represents the average ± S.E.M. of at least three independent experiments. Error bars are present for all data points/columns on all plots in all figures but may not be visible. For time course experiments, the initial assay solution contained 6 pmol of a Cy3/Cy5-labeled hairpin (**Figure S1**) and the initial E_FRET_ trace was recorded. The recording was paused, excess RPA (90 pmol) was added, and the E_FRET_ trace of the resultant solution was recorded beginning 3 s after the addition of RPA. Finally, the recording was paused, excess unlabeled poly(dT)_70_ ssDNA (2250 pmol) was added, and the E_FRET_ trace of the resultant solution was recorded beginning 3 s after the addition of poly(dT)_70_. Data is plotted as a function of time with time courses adjusted for the time between each addition (RPA, poly(dT)_70_) and the recording of E_FRET_ (Δt = 3 s). For all plots of E_FRET_ Atraces in all figures, each trace represents the average ± S.E.M. of at least three independent experiments. Error bars are present for all time points of all traces on all plots in all figures.

### Primer Extension Assays

All primer extension experiments were performed room temperature (23 ± 2 °C) in 1X Replication Buffer supplemented with 1 mM DTT and 1 mM ATP, and the final ionic strength was adjusted to physiological (200 mM) by the addition of appropriate amounts of KOAc. All reagents, substrate, and protein concentrations listed are final reaction concentrations. For all experiments, PCNA was first loaded onto a Cy5-labeled P/T DNA as follows. A Cy5-labeled P/T DNA (250 nM, **Figure S1**) was preincubated first with NeutrAvidin (1 μM homotetramer) and then RPA (3.75 μM heterotrimer). Then, PCNA (250 nM homotrimer), ATP (1 mM), and RFC (250 nM heteropentamer) were added in succession and the resultant mixture was incubated to load PCNA. For time course experiments, dNTP incorporation (i.e., primer extension) was initiated by the successive addition of dNTPs (46 μM dATP, 9.7 μM dGTP, 48 μM dCTP, 67 μM dTTP, unless indicated otherwise) and pol δ (25 nM heterotetramer). The concentration of each dNTP utilized was within the physiological range observed in dividing human cells (39). At variable times, aliquots of the reaction were removed, and quenched with 62.5 mM EDTA, pH 8.0, 2 M Urea, 50 % formamide supplemented with 0.01 % (wt/vol) tracking fluorophores. For single time point experiments, pol δ (25 nM heterotetramer) was added after PCNA loading, the resultant mixture was incubated and divided into 5 equal volume aliquots. Primer extension was then initiated for each aliquot by the simultaneous addition of dNTPs and a given concentration of poly(dT)_70_ (0 – 1.875 μM). After ∼1.5 min, the reaction was quenched as described above.

All primer extension products were resolved on 8 % sequencing gels via denaturing PAGE. Before loading onto a gel, quenched samples were heated at 95 °C for 5 min and immediately chilled in ice water for 5 min. Gel images were obtained on a Typhoon Model 9410 fluorescence imager (GE Healthcare). The fluorescence intensity of each DNA band within a given lane on a gel was quantified with ImageQuant (GE Healthcare) and then converted to concentration by first dividing its intensity by the sum of the intensities of all DNA bands present in the respective lane and then multiplying the resultant fraction by the concentration of P/T DNA (250 nM). The percent encounter for a given DNA template sequence (% Encounter) is the percentage of pol δ holoenzymes that initiate DNA incorporation and encounter the respective DNA template sequence. Within a given lane on a gel, this was calculated by first dividing the concentration of all primer extension products resulting from dNTP incorporation through at least the template nt immediately upstream (i.e., 3’) of the respective DNA template sequence (i.e., Encounter Products) by the concentration of all primer extension products and then multiplying the resultant fraction by 100%. The percent transit for a given DNA sequence (% Transit) is the percentage of progressing pol δ holoenzymes that encounter a given DNA template sequence and continue dNTP incorporation completely through the respective DNA template sequence. Within a given lane on a gel, this was calculated by first dividing the concentration of all primer extension products resulting from dNTP incorporation through at least the last nt of the respective DNA template sequence (i.e., Transit Products) by the concentration of Encounter Products (described above) and then multiplying the resultant fraction by 100%. Only data points within 20% of the reaction progress (based on the accumulation of primer extension products) are displayed in the gel images and considered for further analyses. Within this time frame (i.e., linear/steady state phase) the parameters discussed above remain constant with incubation time (40). Importantly, for all initial dNTP concentrations utilized in the present study, primer extension reduced the concentration of each dNTP ≤ 5% over the first 20% of the reaction progress. For all plots in all figures, each data point/column represents the average ± S.E.M. of at least three independent experiments. Error bars are present for all data points/columns on all plots in all figures but may not be visible.

## RESULTS

### The conformations of [AT/TA]_25_ microsatellite repeats are dependent on RPA availability

First, we utilized FRET to investigate the structural topology of [AT/TA] microsatellite repeat sequences within the human genome. FRA3B-AT3 is a 90 nt forward DNA sequence that contains a 3’-[AT]_25_-5’ sequence flanked on the 5’ and 3’ ends by 6 nt and 34 nt, respectively (**Table 1**). FRA3B-AT3-RC is the reverse complement of FRA3B-AT3 and both DNA sequences are predicted to form a hairpin with a 4 nt loop and a perfectly matched 24 A:T base pair (bp) stem that is closed by a G:C bp (32-35). Both DNA sequences are labeled with an internal FRET pair comprised of a cyanine 3 (Cy3) and a cyanine 5 (Cy5) fluorophore (**Figure 1A, Figure S1**). The location of the FRET pair was selected to account for multiple DNA conformations. Internal Cy3/Cy5 FRET pairs have minimal effects on the local thermodynamic stability of double-stranded DNA (dsDNA) constructs (41). On a native PAGE gel, both Cy3/Cy5-labeled DNA sequences were observed predominantly as a single band that migrates slightly faster than the control DNA (**Figure 1B**), indicating that both DNA sequences exist predominantly as their predicted hairpin (depicted in **Figure 1A**), rather than a homodimer that would be 52 bp in size and migrate much slower than the control DNA.

The affinity of human RPA for ssDNA is exceptionally high (∼ nM to pM range) and exceeds that of any ssDNA-binding protein/protein complex that has been identified in human cells. During scheduled DNA replication in S-phase, RPA immediately engages DNA templates exposed within the replisome, extending the bound sequence into a linear conformation and increasing its bending 2 – 3-fold (30,42-44). This prevents the engaged DNA sequence from collapsing into stable, intramolecular secondary structures, such as G quadruplexes (44-47). To investigate the effect of RPA on the structure of [AT/TA] microsatellite repeats, each Cy3/Cy5-labeled DNA sequence described above was titrated with RPA and the FRET signal (i.e., E_FRET_) observed at equilibrium after each addition of RPA was monitored. Here, the observed E_FRET_ values directly report on the distance between the internal cyanines at equilibrium and, hence, the structural topology of the [AT/TA]_25_ sequence.

In the absence of RPA, high E_FRET_ values were observed at equilibrium for each Cy3/Cy5-labeled DNA sequence (**Figure 1C**), as expected for the hairpin conformation where the internal cyanines are juxtaposed in the duplex region (as depicted in **Figure 1A**) and within 20 Å of each other. Differences in the equilibrium E_FRET_ values observed in the absence of RPA likely reflect differential effects of the 33 nt overhangs on the fluorescence intensities of the cyanines due to distinct interactions of these overhangs with the hairpin stem (48). Upon titration with RPA, the equilibrium E_FRET_ values decrease incrementally until an [RPA]:[DNA] ratio of ∼12.5 and then persist at a minimum E_FRET_ value that is identical to the background E_FRET_ for the fully-unwound (i.e., linear) conformation (**Figure S2**) where the internal cyanines are separated by 18 nt of linear ssDNA, which spans a distance (113.4 – 121.7 Å) significantly greater than the Cy3/Cy5 FRET limit (∼ 100 Å) (49). This indicates that saturation of the Cy3/Cy5-labeled DNA sequences with RPA fully unwinds the hairpins and stabilizes the [AT/TA]_25_ sequences in linear conformations. Thus, FRA3B-AT3 and FRA3B-AT3-RC are each in; 1) a hairpin conformation when devoid of RPA; 2) a linear conformation when saturated with RPA.

Next, we monitored E_FRET_ of the Cy3/Cy5-labeled DNA sequences over time to investigate the interplay between RPA availability and the structural topology of [AT/TA] microsatellite repeats (**Figure 2A**). First, a Cy3/Cy5-labeled hairpin was incubated in isolation. Next, RPA was added at a 15-fold excess of the respective hairpin concentration to fully unwind the hairpin into a linear conformation (**Figure 1C**). The predominant ssDNA-binding footprint of human RPA is ∼30 nt (45,50-53). Hence, the linear conformation of each Cy3/Cy5-labeled DNA sequence accommodates at least 3 RPA complexes, which form a directional filament (as depicted in **Figure 2A**)(50,51,53-58). Next, poly(dT)_70_ was added at a concentration that is 15-fold greater than the concentration of RPA and 375-fold greater than the concentration of the respective Cy3/Cy5-labeled DNA sequence. Poly(dT) sequences are incapable of adopting secondary structures and exist as a collection of collapsed, completely flexible states (59). Human RPA tightly engages and fully extends poly(dT) sequences (**Figure S2**) (30,43,50-53). Under these conditions, poly(dT)_70_ serves as both a passive and active trap of RPA; any free RPA (in solution) is passively captured by the trap and prevented from binding a Cy3/Cy5-labeled DNA sequence; RPA engaged with a Cy3/Cy5-labeled DNA sequence is actively transferred to the trap via facilitated exchange and prohibited from re-binding to a Cy3/Cy5-labeled DNA sequence (36). For these analyses, E_FRET_ traces were normalized to their respective range to more clearly report on the structural topology of the [AT/TA]_25_ sequences. Specifically, for each Cy3/Cy5-labeled DNA sequence, E_FRET_ values observed in the absence of RPA and poly(dT)_70_ served as the maximum of the range and reflected the hairpin conformation. The background E_FRET_ value (0.270 + 0.00367, **Figure S2**) described above served as the minimum of the range and reflected the fully unwound, linear conformation.

**Figure 2.**
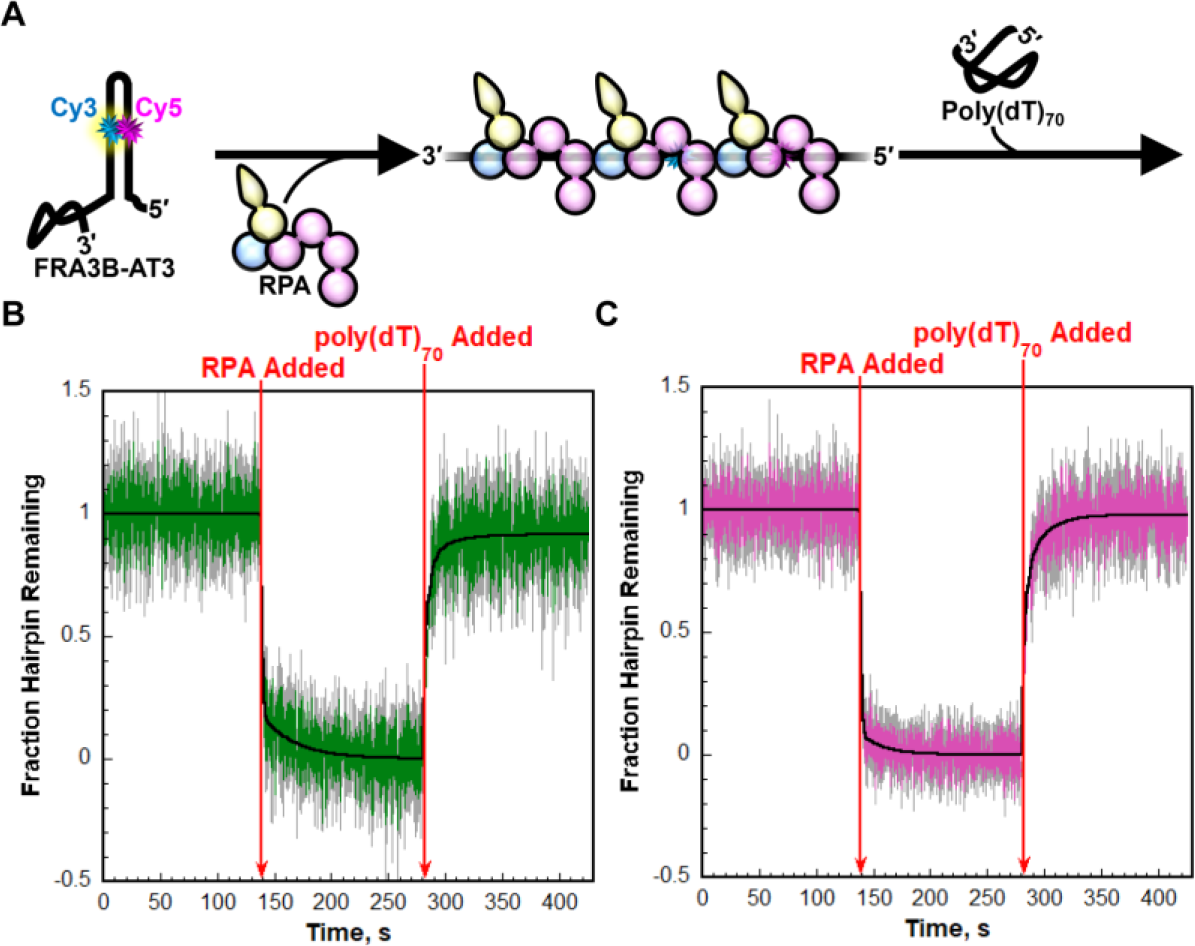
Rapid unwinding of [AT/TA]_25_ hairpins by saturating RPA is completely reversible. (**A**) Schematic cartoon representation of the FRET experiment to monitor the structural dynamics of the FRA3B-AT3 DNA hairpins. As an example, the FRA3B-AT3 DNA hairpin is depicted. (**B** – **C**) Normalized FRET data. Shown in panels **B** and **C** are the normalized E_FRET_ traces for the Cy3/Cy5-labeled FRA3B-AT3 and FRA3B-AT3-RC DNA hairpins, respectively. Each trace represents the average of the normalized traces ± S.E.M. from at least three independent experiments. The times at which RPA and poly(dT)_70_ were added are indicated by red arrows in each panel. The traces observed in the absence of RPA and poly(dT)_70_ are fit to flat lines, traces recorded after the addition of RPA were each fit to a double exponential decay, and traces recorded after the addition of poly(dT)_70_ were each fit to a double exponential rise. Kinetic parameters are reported in **Table 2**.

The normalized E_FRET_ values for each Cy3/Cy5-labeled hairpin rapidly decreased to zero upon addition of RPA (**Figure 2B** and **2C**), indicating that saturating RPA rapidly and completely unwinds the hairpins into linear conformations. For each, unwinding is biphasic and the faster phase (i.e., *k*_obs 1_) accounts for more than ∼80% of the total amplitude (A_T_) and is more than ∼18-fold faster than the slower phase (i.e., *k*_obs 2_) (**Table 2**). Upon addition of poly(dT)_70_, the normalized E_FRET_ values for each Cy3/Cy5-labeled DNA sequence rapidly increased to within experimental error of the respective values observed in the absence of both RPA and poly(dT)_70_, indicating that the linear conformations of the Cy3/Cy5-labeled DNA sequences rapidly re-anneal into their hairpin conformations upon complete depletion of RPA. Thus, rapid unwinding of the [AT/TA]_25_ hairpins by saturating RPA is completely reversible. For each Cy3/Cy5-labeled DNA sequence, re-annealing is biphasic and the faster phase (*k*_obs 1_) accounts for more than ∼70% of the total amplitude (A_T_) and is more than ∼5-fold faster than the slower phase (*k*_obs 2_) (**Table 2**). Together, the results presented in **Figures 1** and **2** suggest that the conformations of the [AT/TA]_25_ microsatellite repeats within the AT3 region of the human FRA3B CFS are dependent upon the availability of RPA.

**Table 2.**
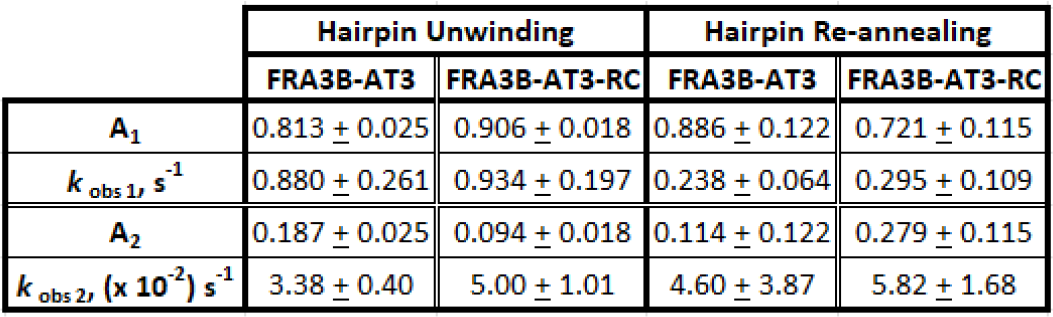
Kinetics parameters of FRA3B-AT3 dynamics

### Stalling of pol δ holoenzymes is not promoted at/within [AT/TA]_25_ microsatellite repeats under native conditions

In humans, lagging strand DNA templates are primarily replicated by pol δ holoenzymes comprised of pol δ and PCNA (24-27). Hence, A+T-rich flexibility regions in lagging strand DNA templates are initially encountered by pol δ holoenzymes during scheduled DNA replication in S-phase. To quantitatively analyze these encounters, we utilized a primer extension assay recently developed by our lab that monitors progression of human pol δ holoenzymes during initial encounters with a DNA template sequence at single nt resolution (40). All experiments are carried out at physiological pH and ionic strength and utilize primer/template (P/T) DNA substrates that mimic nascent P/T junctions on lagging strand templates (**Figure 3**). All P/T DNA substrates are comprised of an identical dsDNA region adjacent to a unique DNA template that is to be replicated by pol δ; either FRA3B-AT3, FRA3B-AT3-RC, or FRA16D-C. Herein, DNA templates are simply referred to as templates. The 29 nt (i.e., 29-mer) primer strand is labeled with a biotin at the 5’ terminus and an internal Cy5 fluorophore 4 nt from the 5’ terminus. The lengths of the dsDNA region (29 bp) and templates (90 nt) support the stepwise assembly of a single pol δ holoenzyme on a P/T junction, as follows (29,36-38,40). First, a P/T DNA substrate is saturated with NeutrAvidin and RPA ([RPA]:[DNA] = 15). NeutrAvidin tightly engages the resident biotin and the resultant complex blocks loaded PCNA from diffusing off the dsDNA end of a P/T DNA substrate. RPA engages and unwinds a template that is to be replicated (**Figure 1C**) and blocks loaded PCNA from diffusing off the ssDNA end of a P/T DNA substrate. Next, PCNA is loaded onto a P/T DNA substrate by RFC and stabilized by RPA and the NeutrAvidin/biotin complex (37,38). In the primer extension assay, dNTP incorporation is initiated by the sequential addition dNTPs and limiting pol δ (**Figure 4A**). The concentration of each dNTP utilized was within the physiological range observed in dividing human cells, unless noted otherwise (39). Extension of the Cy5-labeled 29-mer primer (i.e., N = 29) is monitored via denaturing PAGE for ≤ 20 % of the reaction (**Figure 4B**). Under these conditions, once a primer is extended and the associated pol δ subsequently disengages, the probability that the extended primer will be re-utilized is negligible. Rather, the dissociated pol δ engages another previously unused primer. In other words, the observed primer extension products reflect a single cycle of dNTP incorporation by pol δ. In this setup, nearly all dNTP incorporation (> 90 %) observed is carried out by pol δ holoenzymes, rather than pol δ alone, and only pol δ holoenzymes are responsible for dNTP incorporation beyond the 5^th^ dNTP incorporation step (40)Δ Loading and stabilization of PCNA onto a given P/T DNA substrate is not affected by the length, sequence, or orientation of the respective template that is to be replicated (**Figure S3**). Thus, any observed effects on dNTP incorporation by pol δ are not attributable to the amount of PCNA encircling the resident P/T junction. In these assays, the progression of pol δ holoenzymes towards and through a given downstream template sequence were analyzed (see **Materials and Methods**). Specifically, the percentage of pol δ holoenzymes that initiate dNTP incorporation and encounter a given sequence in a template prior to pol δ dissociation is referred to as the “% Encounter.” The percentage of progressing pol δ holoenzymes that encounter a given sequence in a template and replicate the entire sequence (i.e., transit) prior to pol δ dissociation is referred to as the “% Transit.”

For FRA3B-AT3, the hairpin-forming (HPF) sequence spans dNTP incorporation steps *i*_34_ to *i*_85_ (**Figure 3**). Encounter of the HPF sequence occurs upon the 33^rd^ dNTP incorporation step (*i*_33_) and this indicated by primer extension products ≥ 62 nt in length (N + 33 = 62) (**Figure 4B**). These products are referred to as “HPF Encounter Products”. Transit of the HPF sequence occurs upon the 85^th^ dNTP incorporation step (*i*_85_) (**Figure 3)** and this is indicated by primer extension products ≥ 114 nt in length (N + 85 = 114) (**Figure 4B**). These products are referred to as “HPF Transit Products”. In time-dependent analyses of the HPF Encounter and HPF Transit Products (**Figure 4C**), both emanated from the origin and increased linearly. The absence of a lag time in the appearance of both products indicates that, upon initiation of dNTP incorporation, the HPF sequence is encountered and transited essentially instantaneously. As shown in **Figure 4D**, the % Encounter and % Transit of the HPF sequence (i.e., % HPF Encounter and % HPF Transit) are 11.42 ± 1.41 % and 52.78 ± 0.957 %, respectively. The latter indicates that more than half of the pol δ holoenzymes that encounter the HPF sequence transit the sequence prior to pol δ dissociation.

**Figure 3.**
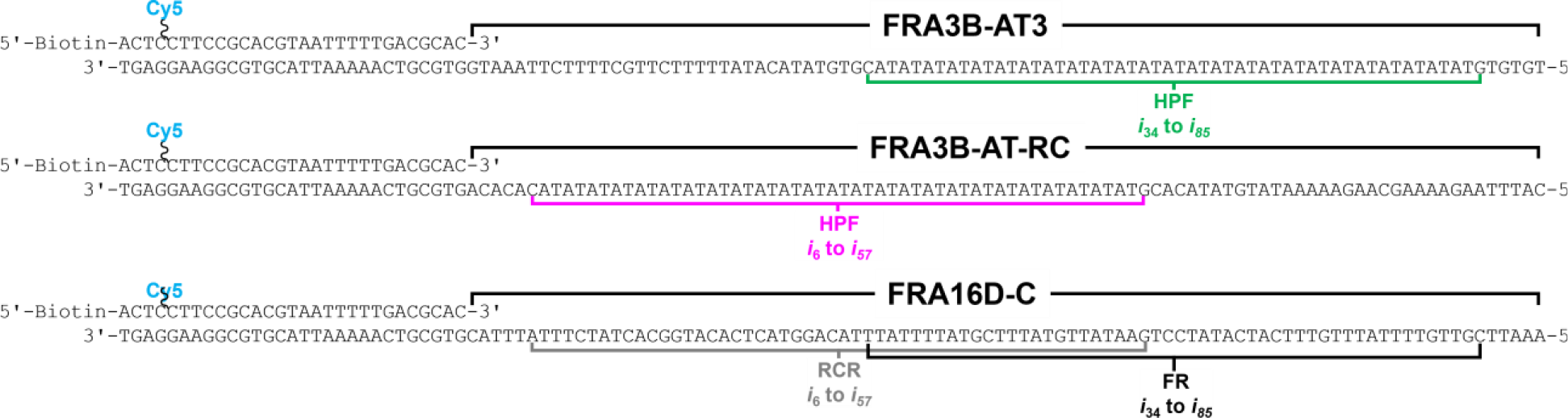
P/T DNA substrates utilized in primer extension assays. Templates to be replicated by pol δ (FRA3B-AT3, FRA3B-AT3-RC, FRA16D-C) are indicated. The HPF sequence in FRA3B-AT3 and FRA3B-AT3-RC and the FR and RCR sequences in FRA16D-C are indicated. dNTP incorporation steps (*i*) within each of these sequences are indicated.

**Figure 4.**
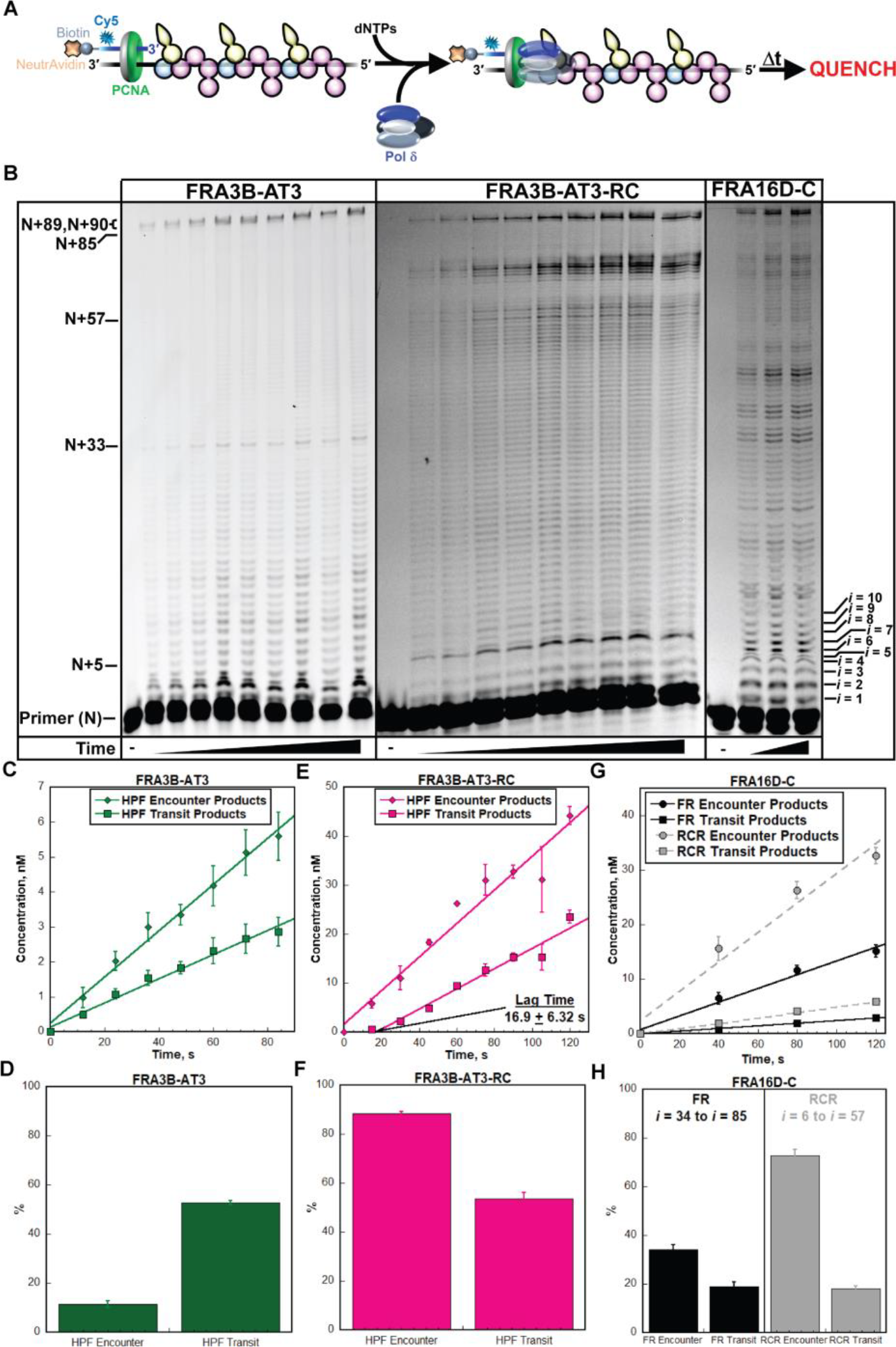
Stalling of pol δ holoenzymes is not promoted at/within [AT/TA]_25_ sequences under native conditions. (**A**) Schematic cartoon representation of the primer extension assay. (**B**) Denaturing sequencing gel of primer extension products observed for FRA3B-AT3, FRA3B-AT3-RC, and FRA16D-C templates. Indicated on the left are the sizes of the primer (N), the minimum primer extension products indicating that a reference or the HPF sequence is encountered or replicated and the full-length products (N+89, N+90). dNTP incorporation step (*i*) for each primer extension product up to 39 nt in length (*i*_10_) is on indicated on the right. (**C** - **H**) Data. Encounter and Transit Products of the HPF sequence in the FRA3B-AT3 and FRAD3B-AT3-RC templates and the reference sequences (FR, RCR) in the FRA16D-C template are shown in panels **C, E**, and **G**, respectively. The % Encounter and % Transit of the HPF sequence in the FRA3B-AT3 and FRAD3B-AT3-RC templates and the reference sequences (FR, RCR) in the FRA16D-C template are shown in panels **D, F**, and **H**, respectively. Each data point/column represents the average + S.E.M. of at least three independent experiments.

For FRA3B-AT3-RC, the HPF sequence spans dNTP incorporation steps i_6_ to i_57_ (**Figure 3**). In this template, encounter of the HPF sequence occurs upon the 5^th^ dNTP incorporation step (*i*_5_) Hence, HPF Encounter Products are ≥ 34 nt in length (N + 5 = 34) (**Figure 4C**). Transit of the HPF sequence occurs upon the 57^th^ dNTP incorporation step (*i*_57_) (**Figure 3**). Hence, HPF Transit Products are ≥ 86 nt in length (N + 57 = 86) (**Figure 4C**). In time-dependent analyses of the HPF Encounter and HPF Transit Products for this template, only the former emanated from the origin; linear regression of the HPF Transit Products crossed the x-axis (time) at 16.9 ± 6.32 s (**Figure 4E**). The presence of a lag time in only the appearance of HPF Transit Products indicates that upon initiation of dNTP incorporation, the HPF sequence is encountered by progressing pol δ holoenzymes essentially instantaneously, but then dNTP incorporation slows such that a significant amount of time elapses before the HPF sequence is transited. For FRA3B-AT3-RC, the % HPF Encounter is 88.38 ± 1.41 % (**Figure 4F)**, which is significantly higher than that observed for FRA3B-AT3 (11.42 ± 1.41 % in **Figure 4D**). This is expected given that the HPF sequence in FRA3B-AT3 is encountered much further downstream of the P/T junction compared to the HPF sequence in FRA3B-AT3-RC (**Figure 3**) and a small portion of progressing pol δ holoenzymes dissociate at each successive dNTP incorporation step (40,60). The % HPF Transit for FRA3B-AT3-RC is 53.64 ± 2.54 % (**Figure 4F**), which is identical to that observed for FRA3B-AT3 (52.78 ± 0.957 %, **Figure 4D**). Thus, despite the significant differences in transit speed through the HPF sequence in the two orientations, pol δ holoenzyme progression through the HPF sequence is neither inhibited nor promoted in one context over the other.

As a control, we repeated these assays on FRA16D-C (**Figure 3**), which has high A+T content (73.3 %) similar to FRA3B-AT3 and FRA3B-AT3-RC but is devoid of any repeat elements that can form stable intramolecular secondary structures. For comparison, the DNA sequences from *i*_34_ to *i*_85_ and from *i*_6_ to *i*_57_ in FRA16D-C are used as references for the HPF sequence in FRA3B-AT3 and FRA3B-AT3-RC, respectively. Herein, these reference sequences are referred to as FR (<u>F</u>orward <u>R</u>eference, *i*_34_ to *i*_85_) and RCR (<u>R</u>everse <u>C</u>omplement <u>R</u>eference, *i*_6_ to *i*_57_). The A + T content of the FR (76.9% A+T) and RCR (69.2% A+T) sequences are similar. In time-dependent analyses of the Encounter and Transit Products for the FR and RCR sequences, all emanated from the origin and increased linearly (**Figure 4G**). The absence of a lag time for the FR Encounter and FR Transit Products agrees with that observed for the HPF Encounter and HPF Transit Products of FRA3B-AT3 (**Figure 4C**), indicating that progressing pol δ holoenzymes encounter and transit the FR sequence in FRA16D-C and the HPF sequence in FRA3B-AT3 with comparable speeds. The absence of a lag time for the RCR Encounter Products (**Figure 4G**) agrees with that observed for the HPF Encounter Products of FRA3B-AT3-RC (**Figure 4E**), indicating that progressing pol δ holoenzymes encounter the RCR sequence of FRA16D-C and the HPF sequence of FRA3B-AT3-RC with comparable speeds.

However, the absence of a lag time for the RCR Transit Products (**Figure 4G**) contrasts with the significant lag time observed for the HPF Transit Products of FRA3B-AT3-RC. Altogether, the data presented in **Figure 4C, 4E**, and **4G** suggests that the distinct slow speed at which progressing pol δ holoenzymes transit the HPF sequence in FRA3B-AT3-RC is not due to the HPF sequence itself but rather the context in which the HPF sequence resides in FRA3B-AT3-RC (i.e., surrounding DNA sequences, distance from the P/T junction, etc.). As shown in **Figure 4H** for FRA16D-C, the % RCR Encounter is significantly higher than the % FR Encounter.

Again, this is expected given that the FR sequence is significantly further downstream of the P/T junction compared to the RCR sequence and a small portion of progressing pol δ holoenzymes dissociate at each successive dNTP incorporation step (40,60). The % FR Transit (18.79 ± 1.93 %) and % RCR Transit (18.12 ± 1.18 %) for FRA16D-C are identical (**Figure 3H**) and significantly lower than the corresponding % HPF transit values observed for FRA3B-AT3 (52.78 ± 0.957 %, **Figure 4D**) and FRA3B-AT3-RC (53.64 ± 2.54 %, **Figure 4F**). This indicates that progression of pol δ holoenzymes is not inhibited at/within a HPF sequence compared to non-structure forming sequences under native conditions. In other words, stalling of pol δ holoenzymes is not promoted at/within a HPF sequence.

### Depletion of dNTPs slows and stalls the progression of pol δ holoenzymes in general

A common DNA replication stress that leads to CFS instability occurs when the demand for dNTPs exceeds their cellular supply (61). This arises from both endogenous and exogenous sources. For example, oncogene activation may prematurely promote DNA replication and cell proliferation, depleting dNTP pools to levels insufficient to support normal replication and genome stability (10). To model this, we repeated the primer extension assays exactly as described above in **Figure 4** except with a 2-fold lower concentration of each dNTP (referred to herein as 0.5X dNTPs) compared to the initial conditions (referred to herein as 1.0X dNTPs). As shown in **Figure 5A** for the HPF sequence in FRA3B-AT3, the lag times for the HPF Encounter and HPF Transit Products increase from 0.0 at 1.0X dNTPs to 5.48 ± 1.48 s and 12.74 ± 2.37 s at 0.5X dNTPs, respectively. Identical behavior is observed for the FR sequence in FRA16D-C where the lag times for the FR Encounter and FR Transit Products increase from 0.0 at 1.0X dNTPs to 3.59 ± 3.16 s and 18.22 ± 3.84 s at 0.5X dNTPs, respectively (**Figure 5B**). For the HPF sequence in FRA3B-AT3-RC, the lag time for the HPF Encounter Products remains at 0.0 for both dNTP conditions whereas the lag time for the HPF Transit Products increases from 16.92 ± 6.32 s at 1.0X dNTPs to 30.90 ± 5.30 s at 0.5X dNTPs (**Figure 5C**). Identical behavior is observed for the RCR sequence in FRA16D-C where the lag time for the RCR Encounter Products remains at 0.0 for both dNTP conditions and the lag time for the RCR Transit Products increases from 0.0 at 1.0X dNTPs to 16.95 ± 4.06 s at 0.5X dNTPs (**Figure 5B**). Thus, depletion of dNTP pools at constant RPA concentration slows the speed at which progressing pol δ holoenzymes encounter and/or transit template sequences.

**Figure 5.**
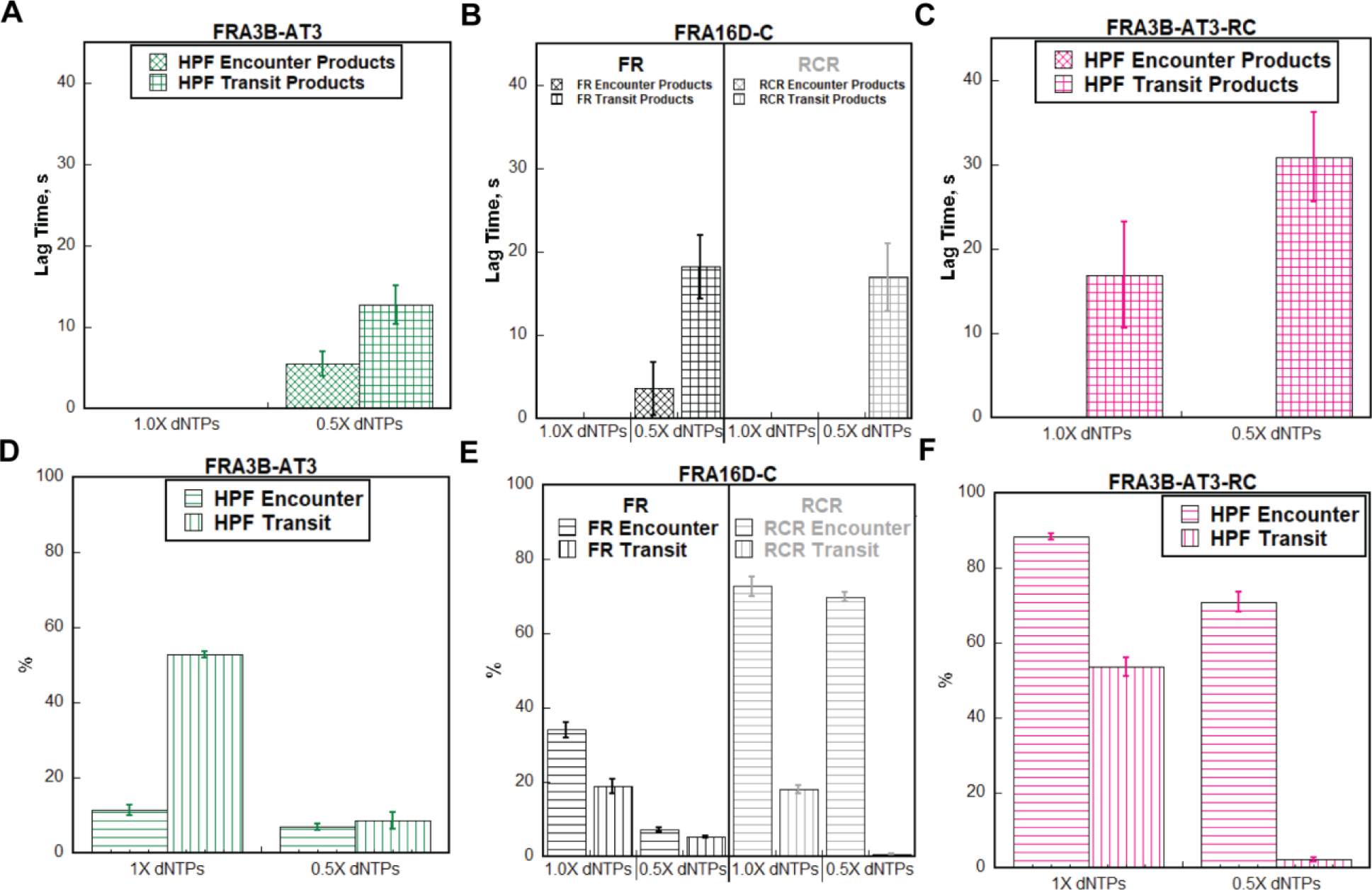
Depletion of dNTP pools at constant RPA concentrations slows and stalls the progression of pol δ holoenzymes in general. Primer extension assays were repeated as depicted in **Figure 4A** - **B** except with a 2-fold lower concentration of each dNTP. Results obtained under these conditions (i.e., 0.5X dNTPs) were analyzed as in **Figure 4C – 4H** and plotted alongside corresponding data obtained under normal conditions (i.e., 1.0X dNTPs). (**A** - **C**) Lag times for the Encounter and Transit Products of the HPF sequence in the FRA3B-AT3 and FRAD3B-AT3-RC templates and the reference sequences (FR and RCR) in the FRA16D-C template. (**D** – **F**) The % Encounter and % Transit of the HPF sequence in the FRA3B-AT3 and FRAD3B-AT3-RC P/T templates and the reference sequences (FR and RCR) in the FRA16D-C templates. Each data column represents the average + S.E.M. of at least three independent experiments.

As shown in **Figure 5D** for the HPF sequence in FRA3B-AT3, the % HPF Encounter is slightly decreased at 0.5X dNTPs by 1.65 ± 0.27-fold and the % HPF Transit is modestly decreased by 6.07 ± 1.60-fold. Similar behaviors are observed for the FR sequence in FRA16D-C (**Figure 5E**) where the % FR Encounter and % FR Transit values are modestly decreased at 0.5X dNTPs by 4.71 ± 0.51 and 3.63 ± 0.42-fold, respectively. For the HPF sequence in FRA3B-AT3-RC, the % HPF Encounter is slightly decreased at 0.5X dNTPs by 1.25 ± 0.05-fold whereas the % HPF Transit is substantially decreased by 24.72 ± 6.23-fold (**Figure 5F**). Similar behaviors are observed for the RCR sequence in FRA16D-C where the % RCR Encounter is essentially maintained at 0.5X dNTPs but the % RCR Transit is substantially decreased by 26.44 ± 4.77-fold, respectively (**Figure 5E**). Thus, in general, depletion of dNTP pools at constant RPA concentration inhibits progression of pol δ holoenzymes towards and/or through template sequences. Collectively, the results presented in **Figure 5** indicate that depletion of dNTP pools at constant RPA concentrations slows and stalls the progression of pol δ holoenzymes in general.

### The effects of RPA on the replication of [AT/TA]_25_ microsatellite repeats is dependent on dNTP concentration

A hallmark of replication stress is the functional uncoupling of enzymatic activities within the replisome such that the rates at which templates are exposed for replication far exceeds the rates at which the exposed templates can be replicated. These uncoupling events expose long stretches of templates that persist for extended periods of time and can arise *in vivo* from endogenous or exogenous sources that restrict DNA replication by depleting dNTP pools (62,63). A previous report revealed that an elevated abundance of exposed templates caused by dNTP depletion also depletes the cellular pool of RPA that is available for subsequent template binding events (i.e. available RPA). This leads to distal stretches of exposed templates that are generated throughout the genome as replication continues (64). To model the effects of this circumstance on the replication of A+T-rich regions, we adapted the primer extension assays described above to monitor dNTP incorporation as conditions transitioned from abundant to depleted dNTP and RPA pools (**Figure 6A**). In short, after PCNA is assembled and stabilized on all P/T junctions, limiting pol δ is added, 5 equal volume aliquots are removed from the resultant mixture, and dNTP incorporation is initiated for each aliquot by the simultaneous addition of dNTPs (either 1.0X or 0.5X) and various concentrations of poly(dT)_70_. The latter rapidly traps RPA while dNTP incorporation is occurring on a given template. The highest concentration of poly(dT)_70_ utilized in these assays completely depletes RPA from a DNA template (**Figure 2**).

**Figure 6.**
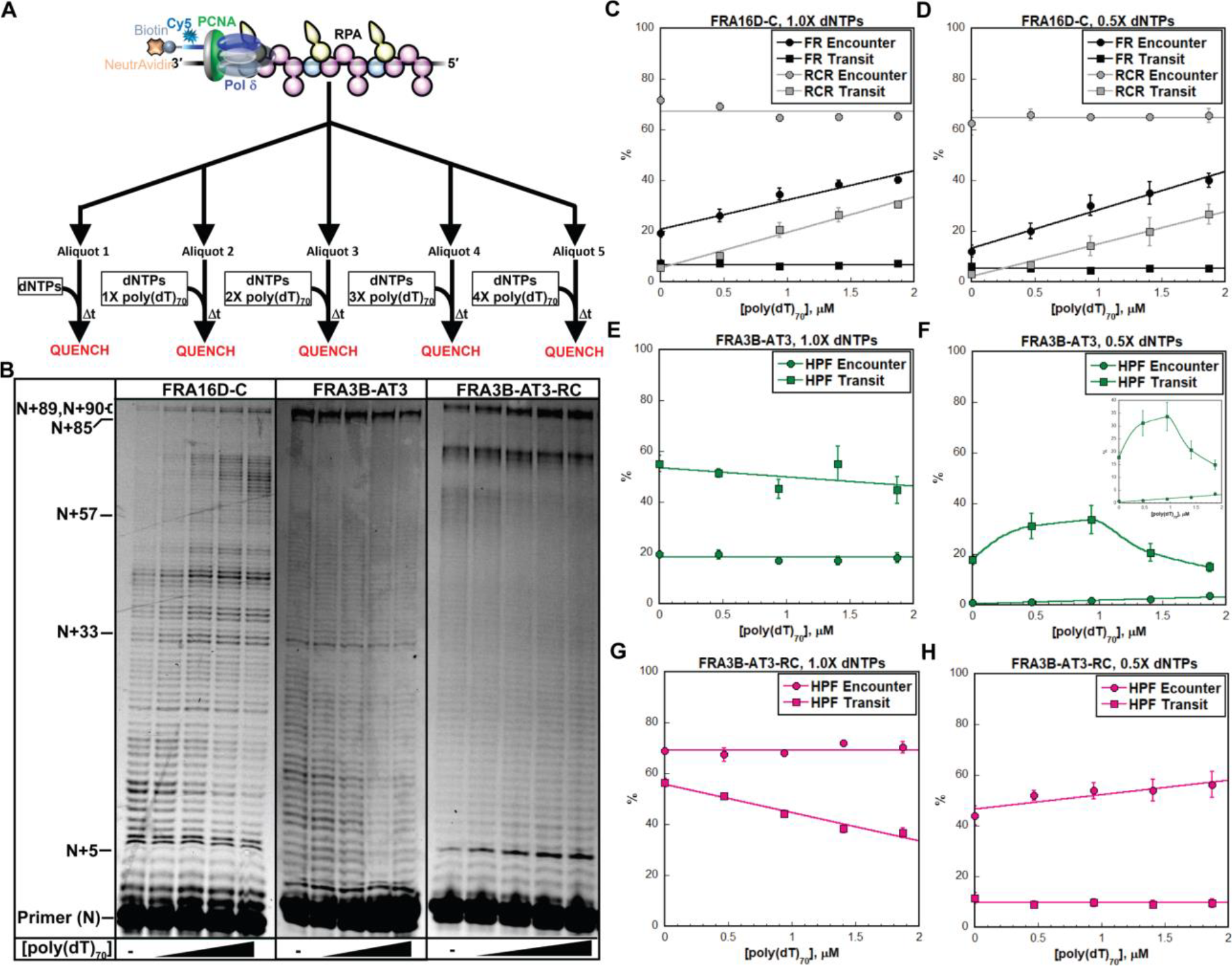
Effects of RPA on the replication of [AT/TA]_25_ sequences is dependent on dNTP concentration. (**A**) Schematic cartoon representation of the experimental procedure. (**B**) 8% denaturing sequencing gel of primer extension products observed for FRA16D-C, FRA3B-AT3, and FRA3B-AT3-RC templates. (**C** - **H**) Primer extension data. The % Encounter and % Transit observed at 1.0X dNTPs for the reference sequences (FR and RCR) in the FRA16D-C template and the HPF sequence in FRA3B-AT3 and FRAD3B-AT3-RC templates are shown in panels **C, E**, and **G**, respectively. The % Encounter and % Transit values observed at 0.5X dNTPs for the reference sequences (FR and RCR) in the FRA16D-C template and the HPF sequence in the FRA3B-AT3 and FRAD3B-AT3-RC P/T templates are shown in panels **D, F**, and **H**, respectively. Each data point represents the average + S.E.M. of at least three independent experiments.

After a period of time (∼1.5 mins) where ≤ 20% of the primer has been extended, each aliquot is quenched, resolved via denaturing PAGE (**Figure 6B**), and analyzed. Because dNTP incorporation for the aliquots emanate from the same pre-assembled pol δ holoenzymes at the same concentration of dNTPs, any differences observed between aliquots are due to the added poly(dT)_70_ and, hence, the extent of RPA depletion. For example, if saturating RPA inhibits pol δ holoenzyme progression, then the % Encounter and/or % Transit for a given template sequence will increase as the concentration of poly(dT)_70_ is increased and RPA is depleted. On the contrary, if saturating RPA promotes pol δ holoenzyme progression, then the % Encounter and/or % Transit for a given template sequence will decrease as the concentration of poly(dT)_70_ is increased and RPA is depleted.

FRA16D-C is devoid of any repeat elements that can form stable intramolecular secondary structures (**Table 1, Figure 3**). At 1.0X dNTPs, the % FR Encounter increased linearly with poly(dT)_70_ concentration (**Figure 6C**), indicating that saturating RPA inhibits progressing pol δ holoenzymes from encountering the FR sequence. However, the % FR Transit was independent of poly(dT)_70_ concentration, indicating that saturating RPA neither promotes nor inhibits pol δ holoenzymes from transiting the FR sequence once they have encountered it. The opposite behavior was observed for the RCR sequence in FRA16D-C. In this case, the % RCR Encounter was independent of poly(dT)_70_ concentration whereas the % RCR Transit increased linearly with poly(dT)_70_ concentration (**Figure 6C**). This indicates that saturating RPA has no effect on progressing pol δ holoenzymes encountering the RCR sequence but inhibits their transit through it. Together, this indicates that saturating RPA promotes the progression of pol δ holoenzymes through the RCR sequence to the FR sequence. Interestingly, the trends observed for the % Encounter and % Transit for both the FR and RCR sequences at 1.0X dNTPs were maintained at 0.5X dNTPs (**Figure 6D**). Thus, the observed effects of RPA on replication of the FR and RCR sequences in the FRA16D-C are independent of dNTP concentration.

FRA3B-AT3 and its reverse complement, FRA3B-AT3-RC, each contain the 3’-[AT]_25_-5’ HPF sequence (**Table 1, Figure 3**). For FRA3B-AT3, the % HPF Encounter was independent of poly(dT)_70_ concentration at 1.0X dNTPs but the % HPF Transit decreased linearly with a shallow but visible slope (**Figure 6E)**. Similar results were observed for the HPF sequence in FRA3B-AT3-RC (**Figure 6G**), albeit the linear decrease in the % HPF transit was much more pronounced. Together, this indicates that saturating RPA has no effect on progressing pol δ holoenzymes encountering the HPF sequence but promotes their transit through it at 1.0X dNTPs to various extents. Interestingly, the trends observed at 1.0X for the HPF sequence in FRA3B-AT3 and FRA3B-AT3-RC were not maintained at 0.5X dNTPs and also partially diverged from one another. Specifically, the % HPF Encounter for both templates at 0.5X dNTPs increased linearly with poly(dT)_70_ (**Figure 6F** and **6H**), in contrast to that observed at 1.0X dNTPs (**Figure 6E 6G**). This indicates that saturating RPA inhibits progressing pol δ holoenzymes from encountering the HPF sequence in each template at 0.5X dNTPs. For FRA3B-AT3, the % HPF Transit at 0.5X dNTPs initially increased with poly(dT)_70_ concentration but then decreased to below the initial value as the concentration of poly(dT)_70_ increased further (**Figure 6F**). This contrasts that observed at 1.0X dNTPs (**Figure 6E**) and reveals that both saturating and completely depleted RPA pools inhibit progressing pol δ holoenzymes from transiting the HPF sequence in FRA3BAT3 at 0.5X dNTPs. For FRA3B-AT3-RC, the % HPF Transit at 0.5X dNTPs was independent of poly(dT)_70_ concentration (**Figure 6H**), indicating that neither saturating RPA nor depleted RPA pools effect transit of the HPF sequence by pol δ holoenzymes. This contrasts that observed at 1.0X dNTPs for FRA3B-AT3-RC (**Figure 6G**) as well as that observed for FRA3B-AT3 at 0.5X dNTPs (**Figure 6F**). Altogether, the results presented in **Figure 6E** – **6H** indicate that the observed effects of RPA on replication of the HPF sequence in FRA3B-AT3 and FRA3B-AT3-RC are dependent on the concentration of dNTPs.

## Discussion

Several CFSs contain A+T-rich, “high flexibility” regions, including stretches of [AT/TA] microsatellite repeats that have the potential to collapse into hairpin structures when in ssDNA form (4-7). Genome-wide analyses revealed that A+T-rich CFS sequences prone to hairpin formation coincide with chromosomal hotspots for genome instability (8-14). While many factors contribute to CFS instability (15,16), direct evidence exists for DNA replication stalling at/within specific A+T-rich CFS regions (3,13,17-20), particularly expanded [TA]_n_ repeats that form intramolecular secondary structures in human cells under replication stress (21-23). In the present study, we characterized the structures of A+T-rich DNA sequences from human CFSs and their replication by pol δ holoenzymes. The results reveal complexities in the structural dynamics and replication of A+T-rich DNA sequences and provide important insights into how replication stress causes DNA replication stalling at/within [AT/TA] microsatellite repeats.

### Depletion of dNTP pools slows and stalls progression of pol δ holoenzymes in general

A common DNA replication stress that leads to CFS instability occurs when the demand for dNTPs exceeds their cellular supply (10,61). The results presented in **Figure 5** indicate that depleting the concentration of dNTPs (from 1.0X to 0.5X) at constant RPA concentration slows and stalls the progression of pol δ holoenzymes in general. This agrees with the established kinetic mechanism for DNA polymerases, as follows. The rate-limiting step for dNTP incorporation is the actual catalysis of dNTP incorporation, which is described by the rate constant *k*_pol_ and occurs after binding an incoming dNTP (65). Thus, *k*_pol_ encompasses both dNTP binding and dNTP incorporation and, hence, is dependent on the concentration of dNTPs (i.e., *k*_pol,obs_ ≤ *k*_pol_). The affinities of human pol δ for each dNTP (*K*_D,dNTP_) are in the low μM range and comparable to the concentration of each dNTP under physiological conditions. For example, the concentration of dGTP in a dividing human cell is 5.2 ± 4.5 μM and the affinity of human pol δ for dGTP is *K*_D,dGTP_ = 3.6 ± 0.8 μM (39,66-68). Thus, *k*_pol_ for all template nts, and hence pol δ holoenzyme progression, is innately limited under physiological conditions (i.e., *k*_pol,obs_ < *k*_pol_) due to sub-saturating dNTP concentrations. Accordingly, a further decrease in the concentration of each dNTP exacerbates these limitations on *k*_pol_ for all template nts, slowing and stalling the progression of pol δ holoenzymes even further (65).

### Effects of RPA on the progression of pol δ holoenzymes through an A+T-rich, non-structure-forming sequence are independent of dNTP concentration

FRA16D-C is devoid of any repeat elements that can form stable intramolecular secondary structures (**Table 1, Figure 3**) (31). Interestingly, progression of pol δ holoenzymes through the RCR sequence to the FR sequence was promoted by the stepwise depletion of RPA at both 1.0X dNTPs (relatively faster *k*_pol,obs_, **Figure 6C**) and 0.5X dNTPs (relatively slower *k*_pol,obs_, **Figure 6D**). This agrees with previous *in vitro* studies carried out in the absence of RPA where progression of human pol δ holoenzymes through FRA16D-C is not significantly hindered (32). Altogether, this indicates that progression of human pol δ holoenzymes on FRA16D-C is inhibited by saturating RPA, promoted by RPA depletion, and these RPA effects are independent of dNTP concentration.

When ssDNA is engaged by RPA complexes, the resultant RPA filament extends the bound ssDNA into a linear conformation and increases its bending rigidity 2 – 3-fold (42,43,47,50-52,69). During scheduled DNA replication in S-phase, this prevents an exposed template from collapsing into stable, intramolecular secondary structures that inhibit DNA replication (44-47). However, in isolation, non-structure forming template sequences, such as FRA16D-C, behave as completely flexible polymers at physiological ionic strength and, hence, exist as a collection of collapsed, highly dynamic conformations in which the constituent template nts are readily accessible (49,59,70). Hence, the ability of RPA to unwind and/or prevent inhibitory secondary structures offers no benefit to the progression of pol δ holoenzymes through exposed templates that cannot form these structures. Interactions of RPA complexes with exposed templates are, however, competitive with those of a pol δ holoenzyme and, hence, restrict the access of a pol δ holoenzyme to the underlying sequence. Specifically, a progressing pol δ holoenzyme incorporates dNTPs opposite template nts as they are transiently released from an RPA complex. Accordingly, for an exposed template engaged by RPA, replication of each constituent nt is rate-limited either by dNTP incorporation (i.e., *k*_pol,obs_) or by the release of the template nt from an RPA complex, whichever is slower. Thus, we propose that under abundant RPA conditions, saturating RPA inhibits progression of pol δ holoenzymes through non-structure forming sequences, such as FRA16D-C, by restricting access to the underlying sequence. Here, the RPA-engaged template is replicated through multiple pol δ binding encounters. Under depleted RPA conditions, the aforementioned restriction is relieved, promoting progression of pol δ holoenzymes through these freely flexible sequences.

### Effects of RPA on the progression of pol δ holoenzymes through an A+T-rich, HPF sequence are dependent on dNTP concentration

The A+T-rich template FRA3B-AT3 and its reverse complement, FRA3B-AT3-RC, each contain a 3’-[AT]_25_-5’ HPF sequence (**Table 1, Figure 3**) that is stabilized in a linear conformation when saturated with RPA and collapsed into a hairpin structure when completely devoid of RPA (**Figures 1** and **2**). When RPA and dNTPs are abundant (i.e., 1.0X dNTPs), more than 50 % of progressing pol δ holoenzymes that encounter the HPF sequence in these templates transit the sequence prior to dissociation (**Figure 4D** and **4F**). Stepwise depletion of RPA at 1.0X dNTPs inhibits progression of pol δ holoenzymes through the HPF sequence in a linear manner (**Figure 6E** and **6G**). This agrees with previous *in vitro* studies carried out in the absence of RPA where progression of pol δ holoenzyme is significantly blocked within the HPF sequence despite elevated dNTP levels, a vast excess of pol δ over the P/T DNA substrate, and long incubation times (32). Altogether, this suggests that under native conditions (i.e., 1.0X dNTPs, abundant RPA), RPA promotes pol δ holoenzyme progression through the HPF sequence by preventing formation of the hairpin structure.

Interestingly, the effects of stepwise RPA depletion on the progression of pol δ holoenzymes through the HPF sequence in FRA3B-AT3 and FRA3B-AT3-RC observed at 1.0X dNTPs (**Figure 6E** and **6G**, relatively faster *k*_pol,obs_) were not maintained at 0.5X dNTPs (**Figure 6F** and **6H**, relatively slower *k*_pol,obs_). Furthermore, the trends observed for each orientation of the HPF sequence at 0.5X dNTPs diverged from one another. Specifically, partial depletion of RPA from FRA3B-AT3 at 0.5X dNTPs promoted progression of pol δ holoenzymes through the HPF sequence but further depletion inhibited progression. Based on the discussion above, this suggests that saturating RPA inhibits pol δ holoenzyme progression through the HPF sequence in FRA3B-AT3 by restricting access to the underlying sequence and sub-saturating RPA promotes progression by preventing the HPF sequence from collapsing into a hairpin structure. Thus, under depleted dNTP conditions, partial depletion of RPA from FRA3B-AT3 stimulates pol δ holoenzyme progression through the HPF sequence by promoting access to the underlying sequence but as depletion of RPA from FRA3B-AT3 advances from partial to complete, the resultant hairpin structure stalls the progression of pol δ holoenzymes through the HPF sequence. For FRA3B-AT3-RC, stepwise depletion of RPA at 0.5X dNTPs had no effect on the progression of pol δ holoenzymes through the HPF sequence (**Figure 6H**). Thus, under depleted dNTP conditions, stalling of pol δ holoenzymes at/within HPF sequence in FRA3B-AT3-RC is independent of RPA interactions with the template and the structure of the HPF sequence.

Altogether, the data presented in **Figure 6E** – **6H** for FRA3B-AT3 and FRA3B-AT3-RC suggests that stalling of pol δ holoenzymes at/within A+T-rich, HPF sequences is highly complex and depends on the sequence context of the HPF sequence, the concentration of dNTPs, and the availability of RPA.

In summary, the present study reveals that the contribution of “high flexibility” regions to the instability of CFSs is highly complex and dependent on the ability of the flexibility region to adopt stable intramolecular secondary DNA structures, the sequence context of the “high flexibility” region, the concentration of dNTPs, the availability of RPA, and likely other cellular variables pertinent to replication such as the abundance of additional accessory factors, activation cellular checkpoints, etc. Further studies are currently underway to further validate this model.

## SUPPLEMENTAL INFORMATION

Supplemental Information includes Supplemental Experimental Methods and Supplemental Figures S1 – S3 and can be found with this article online at ..

## ACKNOWLEDGEMENTS

We would like to thank all members of the Hedglin and Eckert labs for their efforts in reviewing/proofreading the current manuscript. Funding for this research was provided by the National Institutes of Health award number R01 CA237153 to K.A.E. The content is solely the responsibility of the authors and does not necessarily represent the official views of the funders.

## AUTHOR CONTRIBUTIONS

K.G.P and R.L.D. expressed, purified, and characterized all proteins utilized in the present study. K.G.P. performed all experiments and analyzed the data. M.H. designed the experiments and analyzed the data. M.H., K.G.P., and K.A.E wrote the paper.

