## Supporting Information for "Replication of [AT/TA]_25_ microsatellite sequences by human DNA polymerase δ holoenzymes is dependent on dNTP and RPA levels"

**Affiliations**:
[1]Department of Chemistry, The Pennsylvania State University, University Park, PA 16802

[2]Department of Pathology and Laboratory Medicine, The Jake Gittlen Laboratories for Cancer Research, Hershey, PA 17033

### Supplemental Experimental Methods

#### FRET Measurements

All experiments were performed at room temperature (23 ± 2 °C) in 1X Replication Buffer supplemented with 1 mM DTT, and the final ionic strength was adjusted to physiological (200 mM) by the addition of appropriate amounts of KOAc. All measurements were performed in 16.100F-Q-10/Z15 sub-micro fluorometer cells (Starna Cells) and analyzed in a Horiba Scientific Duetta-Bio fluorescence/absorbance spectrometer. Excitation and emission slit widths were each set to 5 nm, unless indicated otherwise. For assays monitoring the extent of RPA-mediated unwinding, a Cy3/Cy5-labeled poly(dT)$_{30}$ ssDNA sequence was titrated with increasing concentrations of RPA and the equilibrium $E_{FRET}$ value at each RPA concentration was calculated exactly as described in the main text. $E_{FRET}$ values for complete unwinding by RPA were then calculated based on a published FRET-based assay (1-3). For assays monitoring the extent of RFC-catalyzed loading of PCNA onto P/T DNA substrates, $E_{FRET}$ values for complete loading of PCNA onto a P/T DNA were calculated based on a published FRET-based assay (3-11). In short, a solution containing Cy3 P/T 33C (110 nM, **Figure S1**), NeutrAvidin (440 nM homotetramer) and 1 mM ATP is first pre-incubated with RPA (330 nM heterotrimer), Cy5-PCNA (100 nM homotrimer) and RFC (100 nM heteropentamer) are then sequentially added, the resultant solution is mixed via pipetting, and $E_{FRET}$ is monitored beginning 10 s after the addition of RFC.  Data is plotted as a function of time after RFC addition with time courses adjusted for the time between the addition of RFC and the recording of $E_{FRET}$ ($\Delta t$ = 10 s).

### Supplemental Figures

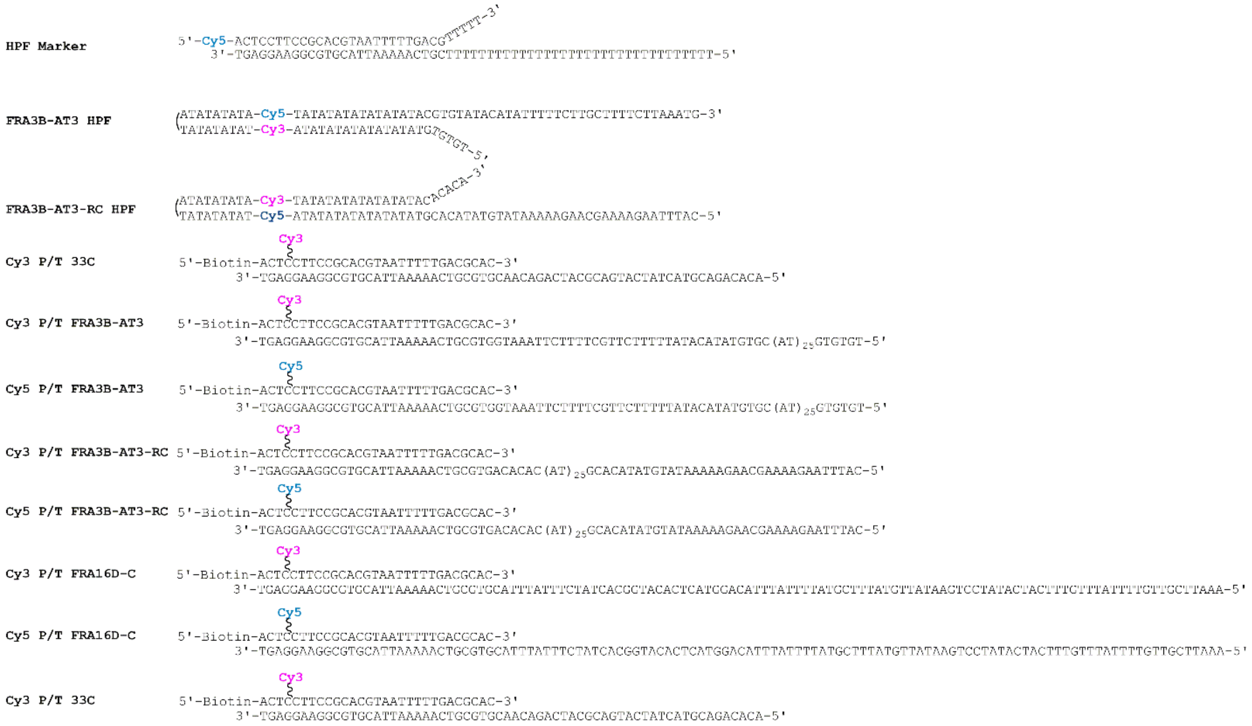

**Figure S1**. DNA utilized in this study. For the P/T DNA substrates, the sequences and lengths (29 bp) of the double-stranded DNA (dsDNA) regions are identical and in agreement with the requirements for assembly of a single PCNA ring onto DNA by RFC (4,7,10). The DNA template sequences to be replicated for these substrates accommodate at least 1 - 3 RPA molecules, depending on the length of the DNA template sequence (12-16). RPA prevents loaded PCNA from sliding off the ssDNA end of the substrate (7). When pre-bound to NeutrAvidin, the biotin attached to the 5’-end of a primer strand prevents loaded PCNA from sliding off the dsDNA end of the substrate.

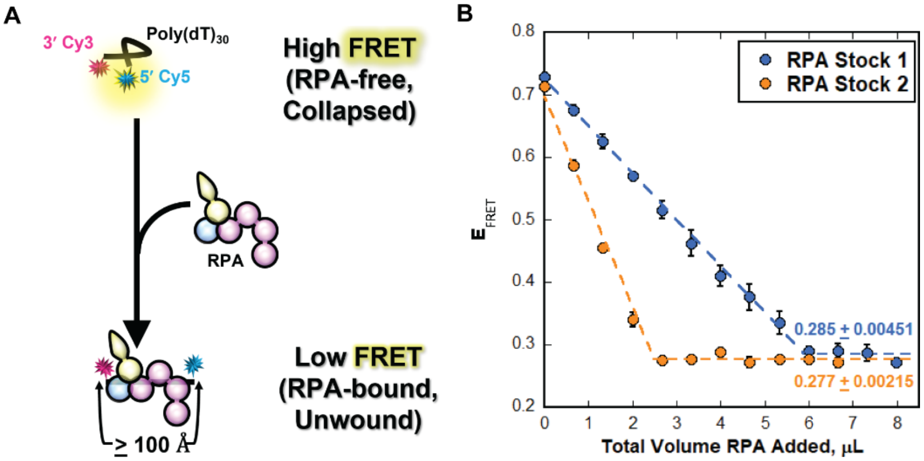


**Figure S2.** FRET-based assay to monitor the extent of RPA-mediated DNA unwinding. The DNA sequence for this assay is a poly(dT)$_{30}$ ssDNA (poly(dT)$_{30}$-FRET) that is terminally labeled with a 3′ Cy3 (FRET donor) and a 5′ Cy5 (FRET acceptor) fluorophore. (**A**) Schematic representation of the FRET-based assay. In the absence of RPA, free DNA (poly(dT)$_{30}$-FRET) forms a compact, flexible structure, bringing the two cyanine fluorophores in close proximity and yielding a robust FRET (i.e., "High FRET" state). At all [RPA]:[DNA] ratios utilized in the assay, the poly(dT)$_{30}$-FRET substrate accommodates 1 RPA complex the 30 nt binding mode (17) and binding of RPA completely unwinds and stretches the engaged ssDNA into a linear configuration and increases its bending 2 – 3 -fold, thereby increasing the Cy3–Cy5 distance to ≥ the Cy3/Cy5 FRET limit (~ 100 Å) and reducing FRET to a background level (i.e., "Low FRET" state) (2,3,18). (**B**) FRET data for RPA titrations carried out with low (RPA Stock 1 = 31.8 ± 1.35 μM) and high (RPA Stock 2 = 111 ± 2.46 μM) concentration RPA stocks. For each, poly(dT)$_{30}$-FRET is titrated with RPA and $E_{FRET}$ is monitored. The observed $E_{FRET}$ [$I_{665}/(I_{665} + I_{563})$] is plotted as a function of the total added volume of RPA and each data point represents the average ± standard error of at least three independent measurements. Under these experimental conditions, binding is stoichiometric and, hence, $E_{FRET}$ decreases linearly until the ssDNA is saturated with RPA (i.e., the equivalence point)(2,3). Each data set is fit to two segment lines; a linear regression with a negative slope and a flat line). $E_{FRET}$ values observed at saturation (indicated) where the ssDNA is completely unwound are calculated from the flat line fits and are essentially identical. The average of the $E_{FRET}$ values observed at saturation for each titration are shown in the plot and are identical. These values were utilized to depict and/or determine the extent of RPA-mediated unwinding of FRA3B-AT3 hairpins in **Figures 1C**, **2B**, and **2C** in the main text.

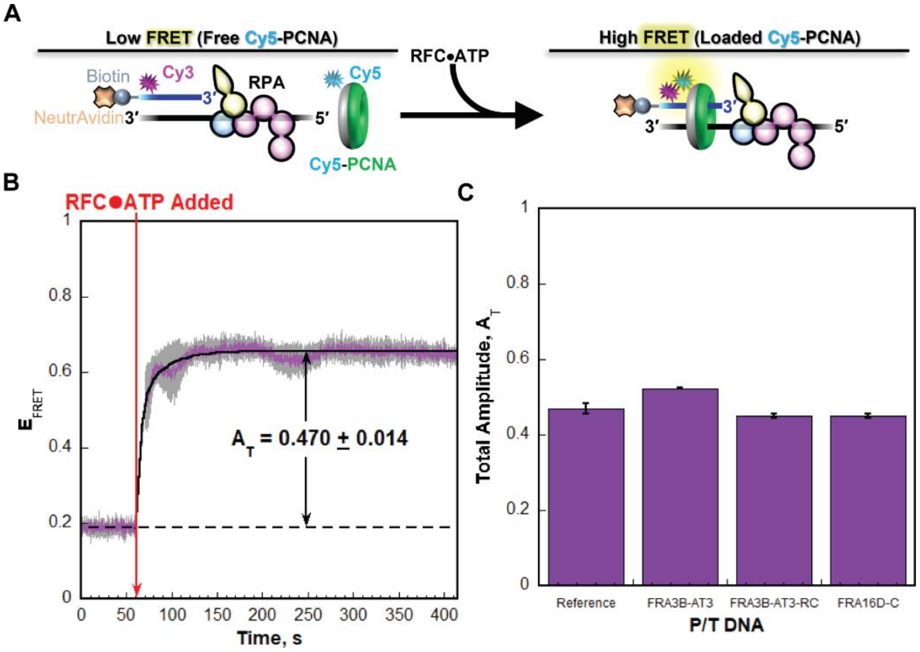


**Figure S3.** FRET-based assay to monitor the extent of RFC-catalyzed loading of PCNA. The DNA substrates for this assay are P/T DNA substrates (**Figure S1**) in which the primer contains a 5′ terminal biotin tag and an internal Cy3 (FRET donor) 4 nt from its 5′ terminus. (**A**) Schematic representation of the FRET-based assay. The Cy3 P/T 33C DNA substrate is shown as an example (**Figure S1**) and contains a non-structure forming, 33 nt DNA template sequence downstream of the P/T junction that accommodates a single RPA molecule. The “back face” of PCNA (shown in grey) is labeled with a Cy5 FRET acceptor. Cy5-PCNA is loaded onto the DNA substrate by the human clamp loader, RFC, such that the Cy5 FRET donor on the “back face” of PCNA faces the Cy3 FRET donor near the 5′ terminus of the primer strand and the “front face” of PCNA (shown in green) is oriented towards the P/T junction where DNA synthesis emanates from. Cy5-PCNA is loaded by RFC in the presence of ATP and excess RPA and $E_{FRET}$ is monitored over time. Under these conditions for this substrate, all Cy5-PCNA is loaded onto a Cy3 P/T 33C by RFC and stabilized by RPA and the biotin/NeutrAvidin blocks that prevent diffusion of PCNA off of the DNA (3-11). (**B**) FRET data for the Cy3 P/T 33C DNA substrate. $E_{FRET}$ trace is the mean of three independent traces with the S.E.M. shown in grey. The time at which the pre-formed RFC•ATP complex added is indicated by a red arrow. The $E_{FRET}$ trace observed prior to the addition of the RFC•ATP complex represents the complete absence of interactions between the Cy3 P/T 33C•RPA complex and Cy5-PCNA and is fit to a flat line that is extrapolated to the axis limits. The $E_{FRET}$ trace observed after the addition of the RFC•ATP complex is fit to a double exponential rise and the overall/total Amplitude ($A_T$) is reported in the graph. For this condition, $A_T$ indicates the change in $E_{FRET}$ observed when all Cy5-PCNA is stably loaded onto the Cy3 P/T 33C DNA substrate. This $A_T$ value (0.4660 ± 0.0145) is utilized as a reference (in panel **C**) to determine the extent of RFC-catalyzed loading of PCNA onto P/T junctions in upstream of the FRA3B-AT3, FRA3B-AT3-RC, or FRA16D-C DNA template sequence (**C**) FRET data for P/T DNA substrates. The assays depicted in panel **A** were carried out and analyzed as described in panel **B**. The overall amplitudes for each P/T DNA substrate utilized in the present study are plotted. All values are in excellent agreement indicating that stable assembly (i.e., loading) of PCNA onto a given P/T junction is not affected by the length, sequence, or orientation of the respective DNA template sequence that is to be replicated.
